# Role of host genetic diversity for susceptibility-to-infection in the evolution of virulence of a plant virus

**DOI:** 10.1101/602201

**Authors:** Rubén González, Anamarija Butković, Santiago F. Elena

## Abstract

Predicting viral emergence is difficult due to the stochastic nature of the underlying processes and the many factors that govern pathogen evolution. Environmental factors affecting the host, the pathogen and the interaction between both are key in emergence. In particular, infectious disease dynamics are affected by spatiotemporal heterogeneity in their environments. A broad knowledge of these factors will allow better estimating where and when viral emergence is more likely to occur. Here we investigate how the population structure for susceptibility-to-infection genes of the plant *Arabidopsis thaliana* shapes the evolution of *Turnip mosaic virus* (TuMV). For doing so we have evolved TuMV lineages in two radically different host population structures: (i) multiple genetically homogeneous *A. thaliana* subpopulations and (ii) a single maximally genetically heterogeneous population. We found faster adaptation of TuMV to homogeneous than to heterogeneous host populations. However, viruses evolved in heterogeneous host populations were more pathogenic and infectious than viruses evolved in the homogeneous population. Furthermore, the viruses evolved in homogeneous populations showed stronger signatures of local specialization than viruses evolved in heterogeneous populations. These results illustrate how the genetic diversity of hosts in an experimental ecosystem favors the evolution of virulence of a pathogen.

## 1. Introduction

Since the term “emerging infectious disease” was coined in the mid-1900s, its definition has evolved (Rosenthal et al. 2015). Woolhouse and Dye (2001) enunciated the most comprehensive definition of an emerging infectious disease as one “*whose incidence is increasing following its first introduction into a new host population or whose incidence is increasing in an existing host population as a result of long-term changes in its underlying epidemiology”.* Engering et al. (2013) rephrased this definition and suggested that emergence events can be classified into three groups: (i) viruses showing up in a novel host, (ii) mutant viral strains displaying novel traits in the same host, and (iii) already known viral diseases spreading out in a new geographic area. Following these definitions, viral emergence is usually associated to cross-species transmission but it can actually occur with or without species jump (Di Giallonardo and Holmes 2015).

The emergence and re-emergence of viruses cause serious threatening to public health (Morens and Fauci 2013) and are responsible for large yield losses in crops that compromise food security (Vurro et al. 2010). Gaining knowledge on the principles that govern viral emergence would allow to predict when and where such events are more likely to happen. Achieving an accurate prediction of the emergence of viral diseases is a challenging task, because emergence is governed by multiple and diverse factors that remain poorly understood or completely unknown. Predictions will become even more difficult under the ongoing climate change that will favor the conditions for development and dispersal of the virus’ vectors (Garrett et al. 2006; Baylis 2017).

The spectrum of disease severity can be attributed to heterogeneity in virus virulence or in host factors; the two are not necessarily independent explanations and they may actually complement and/or interact each other. A problem faced by viruses is that host populations consist of individuals that had different degrees of susceptibility to infection (Schmid-Hempel and Koella 1994; Pfennig 2001). Therefore, adaptive changes improving viral fitness in one host may be selected against, or be neutral, in an alternative one. Genetic variability in susceptibility of hosts and infectiousness of viruses have been well studied in animals and plants (Schmid-Hempel and Koella 1994; Altizer et al. 2006; Brown and Tellier 2011; Anttila et al. 2015; Parrat et al. 2016). Variability within host populations can arise from nonrandom spatial distributions of genotypes: social groups of animals are more closely related to each other than to other members of the population, and plant crops are generally cultivated as genetically homogeneous plots. These situations facilitate virus transmission among genetically similar host genotypes. Spatial structure and local migration predict evolution of less aggressive exploitation in horizontally transmitted parasites (reviewed in Parrat et al. (2016)). For example, Boots and Mealor (2007) observed that *Plodia interpunctella* granulosis virus (PiGV) evolved in spatially-structured lepidopteran host populations become less virulent than the one maintained in well mixed host populations. More recently, Berngruber et al. (2015) showed that a latent λ bacteriophage won competition against a virulent one in a spatially-structured host populations but lost in well mixed populations. In a natural context, it has been observed that virulent *Linum marginale* fungi were more frequent in highly resistant *Melampsora lini* plant populations whereas avirulent pathogens dominated susceptible populations (Thrall and Burdon 2003; Thrall et al. 2012). Similar results were observed in laboratory evolution experiments with the pathosystem *Arabidopsis thaliana – Tobacco etch virus* (TEV): a negative association between plant natural accessions permissiveness to infection and TEV virulence was evolved (Hillung et al. 2014). Finally, host population structure also promotes coexistence of hosts and parasites by creating refugia due to limited dispersal of viruses; increased dispersal makes coexistence less stable (Brockhurst et al. 2006).

The role of host population heterogeneity has also received quite a lot of attention from theoreticians *(e.g.*, Comins et al. 1992; Gandon et al. 1996; Boots and Sasaki 1999, 2002; Gandon and Michalakis 2002; Tellier and Brown 2011), resulting in a number of interesting predictions that in many instances have not been properly tested experimentally. One of the most tantalizing predictions is that in the absence of host heterogeneity, parasites must evolve toward a host exploitation strategy that maximizes transmission with low virulence (Haraguchi and Sasaki 2000; Regoes et al. 2000; Rodríguez and Torres-Sorando 2001; Ganusov et al. 2002; Gandon 2004; Lively 2010; Moreno-Gámez et al. 2013). However, in the context of emerging diseases, right after the spill-over of the new pathogen into the heterogeneous host population and prior to adaptive evolution to take place, Yates et al. (2006) showed that host heterogeneity has a small effect in the probability of establishing the disease. Very recently, Chabas et al. (2018) showed that evolutionary emergence is more likely to occur when the host population contains intermediate levels of resistant hosts, confirming this prediction using different phages and bacterial hosts with different alterations in the CRISPR/Cas immune systems that conferred the cells with different levels of resistance.

In this study we use experimental evolution to explore the effect of host population structure for genes involved in susceptibility to infection in the evolution of infectiousness and virulence of a plant virus; in particular, we want to explore the extent to which host heterogeneity determines the rate of evolution of these two fitness-related traits. According to the above theoretical predictions and experimental observations in other pathosystems, we expect the virus infectiousness and virulence to evolve to lower levels in genetically homogeneous host populations, but at a faster rate, than in maximally genetically diverse host populations. The pathosystem we have studied is composed of *Turnip mosaic virus* (TuMV; genus *Potyvirus*, family *Potyviridae*) as pathogen and six different natural accessions (hereafter ecotypes) of *A. thaliana* that differ in their susceptibility to infection as hosts.

## 2. Methods

### 2.1. Selection of *A. thaliana* ecotypes for evolution experiments

Before we could begin the evolution experiment, we sought to identify a set of *A. thaliana* ecotypes representative of the phenotypic variability in response to TuMV infection (Rubio et al. 2018) observed for a larger collection of ecotypes. Sixteen *A. thaliana* ecotypes were evaluated. Based on the similarity of their phenotypic responses to infection with TuMV (disease intensity over time measured as the area under the disease progression steps *(AUDPS)* curve; see section 2.4 for a definition and explanation) during 14 days post-inoculation (dpi), ecotypes were clustered as shown in Fig. 1 (UPGMA). Six accessions were chosen as representative of the five clusters shown in Fig. 1: Col-0, Ga-0, Gy-0, Oy-0, Ta-0, and Wt-1. A progressive *k*-clustering algorithm confirmed that adding additional clusters did not result in significant improvement in model predictability (five *vs* six clusters partial-*F* test: *F*_1,9_ = 2.841, *p* = 0.058). Ta-0 showed the slowest disease intensity progression (*AUDPS* = 0.490) while Ga-0 the fastest one (*AUDPS* = 6.500); Oy-0 and Wt-1 showed similar progression (*AUDPS* = 1.709).

**Figure 1.**
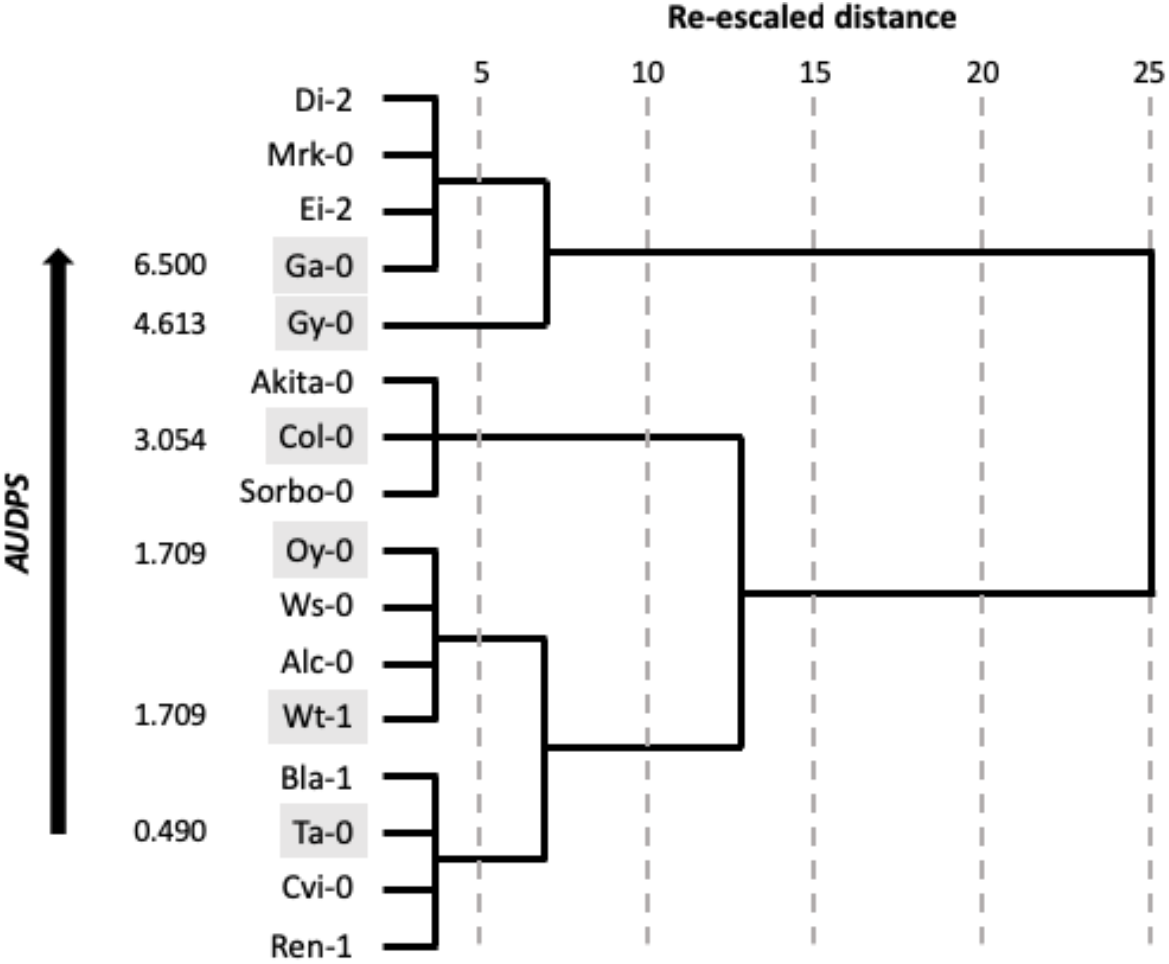
UPGMA clustering of *A. thaliana* ecotypes according to their response to TuMV infection. The six ecotypes selected for the evolution experiment are highlighted in gray. As a measure of virulence, the *AUDPS* of the ancestral TuMV isolate on each selected ecotype is indicated in the left scale.

Furthermore, the six ecotypes reached growth stage 3.5 in the Boyes’ scale (Boyes et al. 2001) at the same time after germination (±1 day), and at that moment they were inoculated. This synchronization ensures that they all were at the same phenological state when inoculated.

In all experiments performed in this study, plants were maintained in a BSL-2 growing chamber at 8 h light: 16 h dark cycles and temperature variation of 24 °C day:20 °C night.

### 2.2. TuMV original isolate and inocula

Infectious saps were obtained from TuMV-infected *Nicotiana benthamiana* plants inoculated with an infectious plasmid containing TuMV genome cDNA (GenBank accession AF530055.2) under the control of the *Cauliflower mosaic virus* 35S promoter. This TuMV sequence variant corresponds to YC5 isolate from calla lily (*Zantedeschia* sp.) (Chen et al. 2003). The same stock of plasmid was used to inoculate two batches of five *N. benthamiana* plants each, with a year of difference. After plants showed visible symptoms of infection, two independent infectious saps (viral stocks) were obtained by grinding infected tissues from *N. benthamiana* plants in a mortar with 10 volumes of grinding buffer (50 mM KH2PO4 pH 7.0, 3% polyethylene glycol 6000).

### 2.3. Experimental evolution

TuMV lineages were evolved during 12 consecutive serial passages as schematized in Fig. 2 in two treatments that differ in the composition of the host population. The first treatment consists in a host population structured in six genetically homogeneous subpopulations (demes). TuMV evolved in this genetically homogeneous host subpopulations. Passages were made by harvesting the symptomatic plants at 14 dpi, preparing infectious sap as described above and inoculating the virus to the next subpopulation (10 plants) of the same ecotype. Three leaves from 21-day old plants were rub-inoculated each with 5 μL of infectious sap and 10% Carborundum (100 mg/mL). One TuMV lineage was evolved on each one of the six ecotypes chosen (Fig. 1). In this treatment, each TuMV lineage in the metapopulation only experienced a particular host genotype along their evolution. In this treatment, we expect evolution to be dominated by rapid local adaptation.

**Figure 2.**
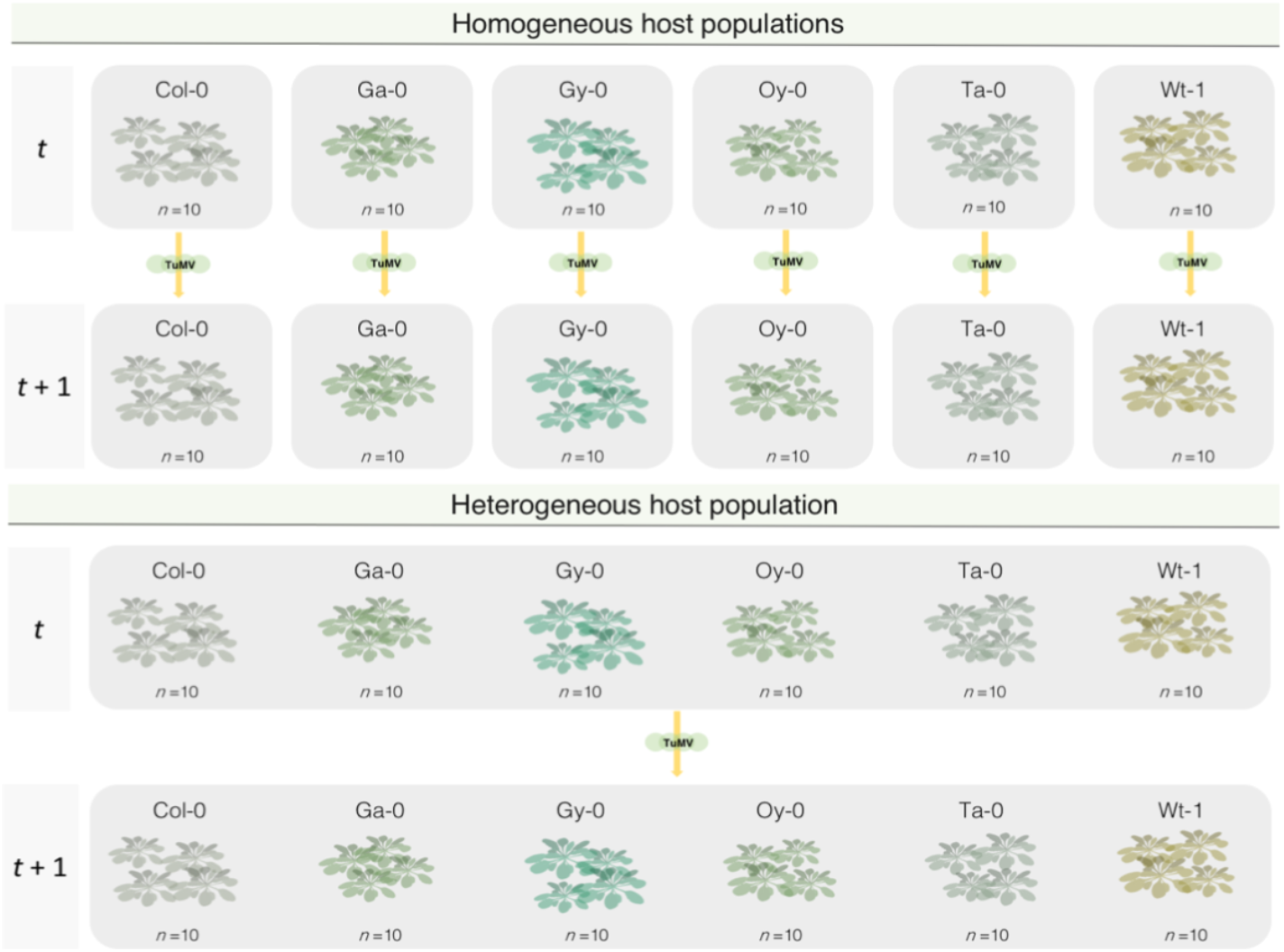
Schematic representation of a passage during the evolution experiments. A total of 12 of such passages were done. See section 2.3 for a detailed description. *A. thaliana* draws were adapted from https://figshare.com/articles/Arabidopsis_Rosette_drawing_steps/4688839.

The second treatment consists of a host population without subpopulation structure composed of 10 plants from each of the six ecotypes. TuMV was thus evolved in a host population with maximal genetic heterogeneity. In this case, the passages were made harvesting all the symptomatic plants of all the ecotypes, pooling the tissues, preparing sap and transmitting it again to 10 plants from each of the ecotypes (Fig. 2). In this treatment, the TuMV evolving population has an equal opportunity of infecting each plant ecotype at each passage. More susceptible ecotypes will contribute more to the next viral generation because more plants are infected and support more generations of replication, while more restrictive ones will contribute in a minor amount to the viral population. In this treatment, we expect a slower adaptation due to fluctuating selection and the evolution of generalist viruses.

This evolution experiment was replicated in two fully independent blocks, hereafter, referred as 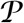 and 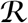, respectively. Each block was initiated with a different viral stock (as described in section 2.2), different light sources (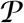 with fluorescent tubes at PAR 100 −150 μmol/m^2^/s and 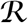 with LED tubes at PAR 90 – 100 μmol/m^2^/s), inoculations were done by two different researchers, and started with more than year of difference in time (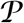 started 12/2/16 and 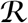 started 1/24/18).

### 2.4. Evaluation of infectiousness and virulence

Upon infection, plants were observed every day and the number of plants showing symptoms was recorded. Infectiousness, *I*, was evaluated as the frequency of plants showing symptoms at 14 dpi out of the 10 plants inoculated with a standardized amount of infectious sap. Using the frequency time-series within the duration of a passage, the *AUDPS* (Simko and Piepho 2012) was evaluated as a proxy to virulence. *AUDPS* represents the speed at which symptoms appear in a population of inoculated plants, and in our case, it is bounded between zero (no plant shows symptoms 14 dpi) and 10 (all plants show symptoms at 1 dpi). In the case of the heterogenous host population treatment, *I* and *AUDPS* were evaluated in each of the ecotypes. *I* data were probit-transformed as 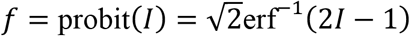 hence the new variable *f* is now Gaussian distributed with mean zero and variance 1. For visualizing infectiousness data in a meaningful scale, *I* will be presented in figures while the *f* transformed data will be used in the statistical analyses described in the section 2.5 below.

At each passage, infection status of each plant was assessed by the presence of symptoms rather than by molecular detection methods. In our extensive experience with the TuMV/*A*. *thaliana* pathosystem, there is an almost one-to-one match between infection and the development of symptoms. Symptoms started with vein bleaching (~5 – 6 dpi) that quickly developed to full leaf chlorosis and necrosis (~10 – 12 dpi). Plants also suffered a developmental arrest, with deformed new leaves, abortion of flowering button and abnormal growth of the caulinar apex.

### 2.5. Data analyses

*AUDPS* and *f* data were fitted to a fully factorial multivariate analysis of covariance model (MANCOVA), in which experimental block (*B*; 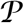 and 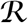), population structure (*D*; homogeneous or heterogeneous) and plant ecotype (*E*) were treated as orthogonal factors and passage (*t*) as a covariable. Main effects and all the interactions between factors and the covariable were incorporated into the model. The full model equation thus reads

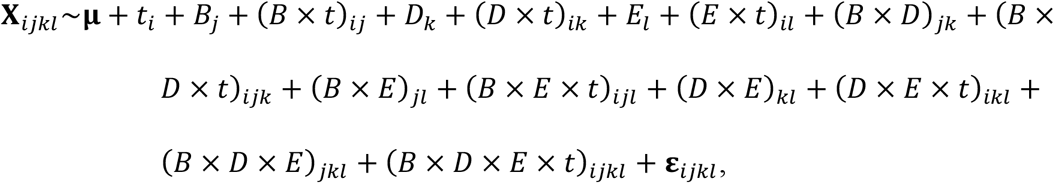

where 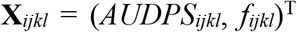 is the vector of phenotypic traits observed at time *i*, experimental block *j*, population structure *k* and ecotype *l*, **μ** represents the vector of phenotypic grand mean values and **ε**_*ijkl*_ stands for the vector of errors. In addition, univariate ANCOVA analyses were performed for *AUDPS* and *f* using the same model equation.

The magnitude of effects was evaluated using the 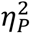 statistic (proportion of total variability in the traits attributable to each factor in the model). Conventionally, values 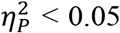 are considered as small, 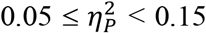 as medium and 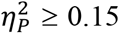 as large effects.

Unless otherwise mentioned, all statistical analyses described in this work were performed using SPSS version 25 software (IBM, Armonk, NY).

### 2.6. Evaluation of rates of phenotypic evolution

*AUDPS* and *I* data obtained for each lineage were fitted to the following first-order autoregressive integrated moving-average, ARIMA(1,0,0), model: 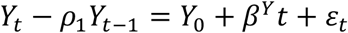, where *Y_k_* represents the variable being analyzed at passage *k*, *ρ*_1_ measures the degree of self-similarity in the time-series data (correlation between values at passages *t* and *t* – 1), *ε_t_* represents the sampling error at passage *t*, and *β^Y^* represents the linear dependency of variable *Y* with passage number, that is, the rate of phenotypic evolution.

To explore the effect of factors *D* and *E* in the rates of *AUDPS* and *I* evolution, the following generalized linear model (GLM) was fitted to the *β* values: 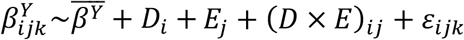, were superscript *Y* again refers to the trait being analyzed, 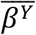 the grand mean value for the rate of evolution of trait *Y* and all other terms are defined in section 2.5. A Normal distribution and identity link function were chosen based on the minimal *BIC* value among competing models. Notice that the error term *ε_ijk_* is obtained from the differences in the estimates from both experimental blocks.

### 2.7. Infection matrices

To analyze the specificity of adaptation of each evolved TuMV lineage, we performed a full cross-infection experiment in which all the 14 evolved lineages were inoculated into 10 plants of all six ecotypes. In the case of the lineages evolved in the genetically heterogeneous host population, the virus isolated at passage 11 was inoculated into 10 additional plants of each of the six ecotypes to separate it into sub-samples. Infection matrices were analyzed using tools borrowed from the field of community ecology to explore whether they show random associations between viral lineages and host genotypes, one-to-one associations, nestedness indicative of a gene-for-gene type of interaction, or modularity (Weitz et al. 2013). The statistical properties of the resulting infection matrices were evaluated using the bipartite version 2.11 package (Dormann et al. 2008) in R version 3.3.2 (R Core Team 2016). Three different summary statistics were evaluated: nestedness (Bascompte et al. 2013), modularity (Newman 2006) and overall specialization *d*’ index (Blüthgen et al. 2006). *d*’ is based in Kullback-Leibler relative entropy, that measures variation within networks and quantifies the degree of specialization of elements within the interaction network. Statistical significance of nestedness and modularity was evaluated using Bascompte et al. (2003) null model.

## 3. Results

### 3.1. Evolutionary dynamics of virulence and infectiousness

Fig. 3 illustrates the evolutionary dynamics of virulence, measured as *AUDPS*, observed on each of the host ecotypes and for both population structures. Likewise, Fig. 4 shows the evolution of infectiousness, *I*, along the passages. In both figures black symbols and lines correspond to the results of experimental block 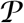 while red symbols and lines correspond to experimental block 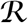. Both variables are significantly correlated (Pearson’s partial correlation coefficient controlling for *B, D, E* and *t*: *r_p_* = 0.640, 306 d.f., *p* < 0.001) and hence multivariate methods were used to analyze the data while increasing the power of the tests. Both datasets were fitted to the MANCOVA model equation described in section 2.5. In all cases, an overall trend to increase virulence and infectivity along time can be observed in the time-series data (Fig. 3 and Fig. 4). This trend is statistically supported by a significant net effect of the passage (covariable *t*) on both phenotypic traits (Table 1; *p* < 0.001).

**Figure 3.**
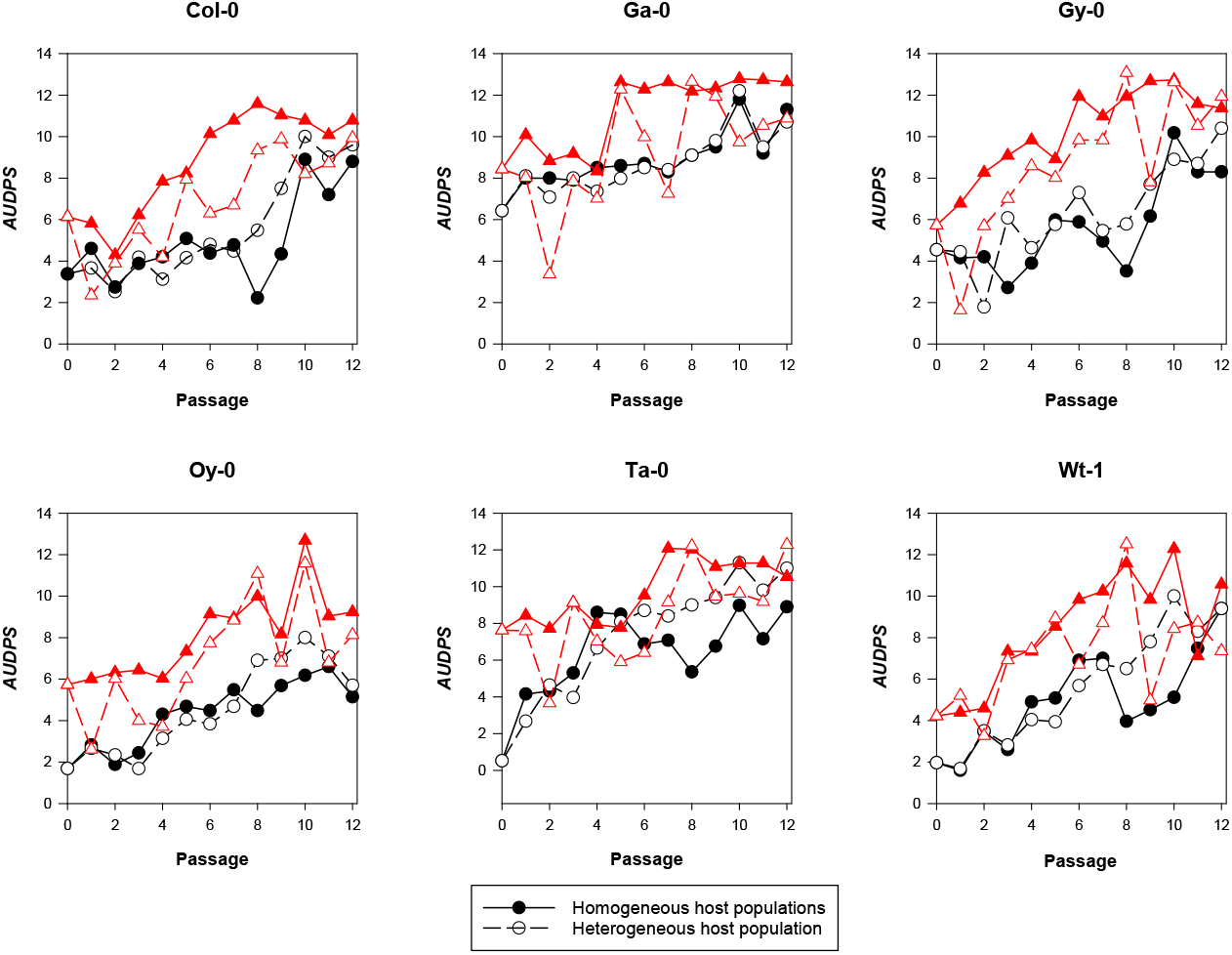
Evolution of virulence *(AUDPS).* Black circles and lines represent the results from experiment 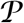 and red triangles and lines from experiment 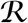. Solid symbols represent evolution in genetically homogeneous host population while open symbols represent evolution in the genetically heterogeneous host populations.

**Table 1.** MANCOVA analysis of the *AUDPS* and *f* data (Fig. 3 and Fig. 4). See section 2.5 for a description of the model equation and parameters. 1 – *β* is the power of the corresponding test.

| Source of variation<sup>1</sup> | Wilk's $\Lambda$ | $F$ | d.f. | $p$ | $\eta_p^2$ | $1 - \beta$ |
| --- | --- | --- | --- | --- | --- | --- |
| *Intersection* ($\mu$) | 0.244 | 406.426 | 2, 263 | < 0.001 | 0.756 | 1 |
| *t* | 0.353 | 241.176 | 2, 263 | < 0.001 | 0.647 | 1 |
| *B* | 0.787 | 35.600 | 2, 263 | < 0.001 | 0.213 | 1 |
| *B*×*t* | 0.911 | 12.805 | 2, 263 | < 0.001 | 0.089 | 0.997 |
| *D* | 0.943 | 7.905 | 2, 263 | < 0.001 | 0.057 | 0.952 |
| *D*×*t* | 0.972 | 3.816 | 2, 263 | 0.023 | 0.028 | 0.690 |
| *E* | 0.713 | 9.678 | 10, 526 | < 0.001 | 0.155 | 1 |
| *E*×*t* | 0.947 | 1.445 | 10, 526 | 0.157 | 0.027 | 0.731 |
| *B*×*D* | 0.993 | 0.969 | 2, 263 | 0.381 | 0.007 | 0.218 |
| *B*×*D*×*t* | 0.985 | 1.979 | 2, 263 | 0.140 | 0.015 | 0.407 |
| *B*×*E* | 0.911 | 2.524 | 10, 526 | 0.006 | 0.046 | 0.954 |
| *B*×*E*×*t* | 0.941 | 1.631 | 10, 526 | 0.094 | 0.030 | 0.794 |
| *D*×*E* | 0.991 | 0.225 | 10, 526 | 0.994 | 0.004 | 0.130 |
| *D*×*E*×*t* | 0.990 | 0.275 | 10, 526 | 0.986 | 0.005 | 0.152 |
| *B*×*D*×*E* | 0.985 | 0.402 | 10, 526 | 0.946 | 0.008 | 0.212 |
| *B*×*D*×*E*×*t* | 0.983 | 0.464 | 10, 526 | 0.913 | 0.016 | 0.316 |

<sup>1</sup>Model factors: passage $\equiv t \in (0, \dots, 12)$ , block $\equiv B \in (\mathcal{P}, \mathcal{R})$ , population structure $\equiv D \in (\text{homogeneous, heterogeneous})$ , host ecotype $\equiv E \in (\text{Col-0, Ga-0, Gy-0, Oy-0, Ta-0, Wt-1})$ .

Next, given the differences in starting inocula, light conditions and experimenter responsible for performing each experimental block, we expect *a priori* to find significant block effects, either net or in combination with other factors in the model. Table 1 shows that experimental block has a net (*B*) effect on the phenotypic vector (*p* < 0.001). This effect changes along evolution passages (*B* × *t*) (*p* < 0.001) and depends on the particular ecotype (*B* × *E*) (*p* = 0.006). The observed net block effect is of large magnitude 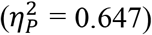, while the effect of its interaction with passage number 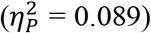 and ecotype 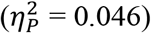 can be considered of small-medium size.

Most interestingly, the population structure in which the TuMV lineages had evolved has a highly significant net (*D*) effect on the phenotypic vector (Table 1; *p* < 0.001), though it is of small-medium size in the multivariate analysis 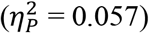, and of similar effect for both *AUDPS* and *f* (univariate ANCOVAs shown in Supplementary Table 1; 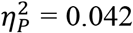 and 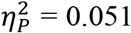, respectively). Taking a look to Fig. 3 and Fig. 4, this effect comes from a pattern (consistent across both experiments) that *AUDPS* and *I* values are usually larger for the lineages evolving in the homogeneous population during the early passages of evolution than for those evolved in the heterogeneous populations (solid symbols are above open ones). Indeed, on average *AUDPS* was 7.42% larger for TuMV lineages evolved in heterogeneous than in homogeneous populations. Likewise, *f* was 12.25% higher for TuMV lineages evolved in the heterogeneous plant population than in the genetically homogeneous ones. Furthermore, the effect of population structure changed along passages (*D* × *t*) (Table 1; *p* = 0.023), though this effect is of small size in the multivariate analysis 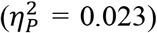 and also in the univariate ones (Supplementary Table 1; 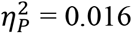 and 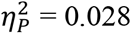, respectively).

**Figure 4.**
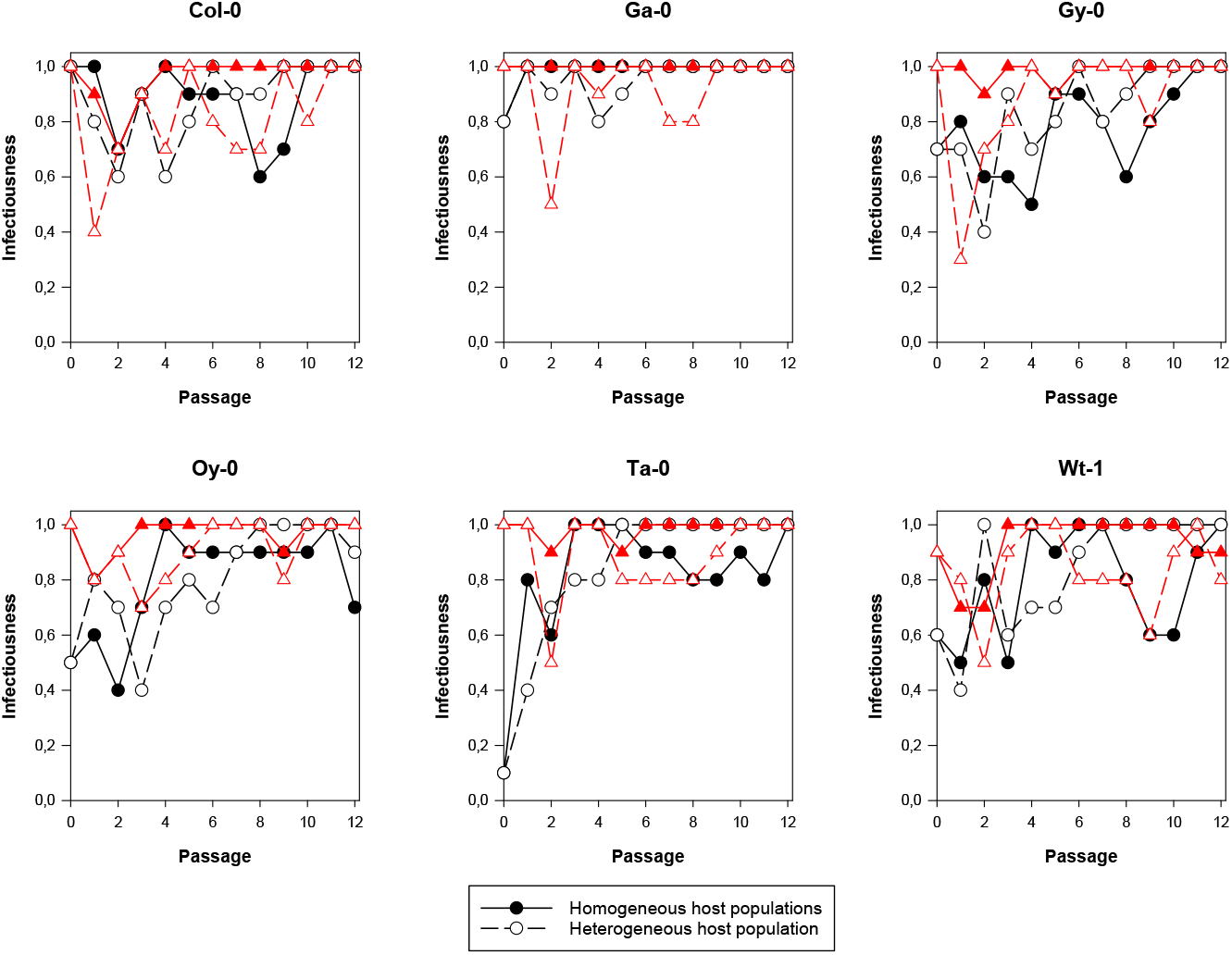
Evolution of infectiousness (*I*). Black circles and lines represent the results from experiment 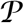 and red triangles and lines from experiment 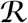. Solid symbols represent evolution in genetically homogeneous host population while open symbols represent evolution in the genetically heterogeneous host populations. Notice that infectiousness data were probit-transformed for statistical analyses.

The host ecotype in which lineages evolved has a highly significant effect on the magnitude of the phenotypic vector (Table 1; *p* < 0.001), the size of this effect being large 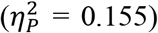. Indeed, the univariate analyses show that the effect is much larger for *AUDPS* than for *f* (Supplementary Table 1; 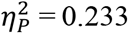 and 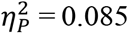, respectively). The TuMV evolved in Ga-0 was the most virulent (*AUDPS* = 9.492), followed by viruses evolved in Ta-0 and Gy-0, while the less virulent infection corresponds to a homogenous group formed by TuMV lineages evolved in Col-0, Wt-1 and Oy-0 (*AUDPS* in the range 5.938 – 6.525) (sequential Bonferroni’s *post hoc* test, *p* ≤ 0.017). This ranking of ecotypes according to virulence slightly differs from the ranking observed for the ancestral TuMV isolate (Fig. 1). Likewise, the virus evolved in Ga-0 was also the most infectious (*f* = 1.264), followed by lineages evolved in Col-0 and Ta-0. The less infectious viruses were those evolved in Wt-1, Oy-0 and Gy-0 (*f* in the range 0.905 – 0.972).

Therefore, we conclude in this first section that the presence of maximal host genetic diversity for genes involved in susceptibility to infection selects for more virulent and infectious viruses. For both phenotypic traits studied, this effect depends on the presence of particular ecotypes.

### 3.2. Rates of phenotypic evolution

In the previous section we found that a net effect of passage on the virulence-related traits. More interestingly, this effect depended on the population structure (*D*) and was consistent across both experimental blocks (non-significant *B* × *D* × *t* effect; Table 1), which allow us to treat the estimates of rates of evolution from each block as replicates in a GLM analysis. Next, we sought to get a better understanding of the effect of population structure for susceptibility to infection on the rates of phenotypic evolution. To do so, we have estimated evolution rates as described in section 2.6. Summary statistics for the ARIMA(1,0,0) model fitting are shown in Supplementary Table 2. Fig. 5 compares the estimated rates of evolution for *AUDPS* and *I* for both population structures analyzed. Rates of evolution were fitted to the GLM described in section 2.6. The results from these analyses are shown in Table 2. Firstly, lets focus in the rate of *AUDPS* evolution. In all cases except for the lineages evolved in Wt-1, the rate of evolution for lineages evolved in the homogeneous host populations is larger than for the corresponding lineages evolved in the heterogeneous host population (Fig. 5A). This trend is statistically significant (Table 2; *p* = 0.015) and the size of effect must be considered as large 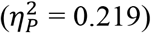.

**Figure 5.**
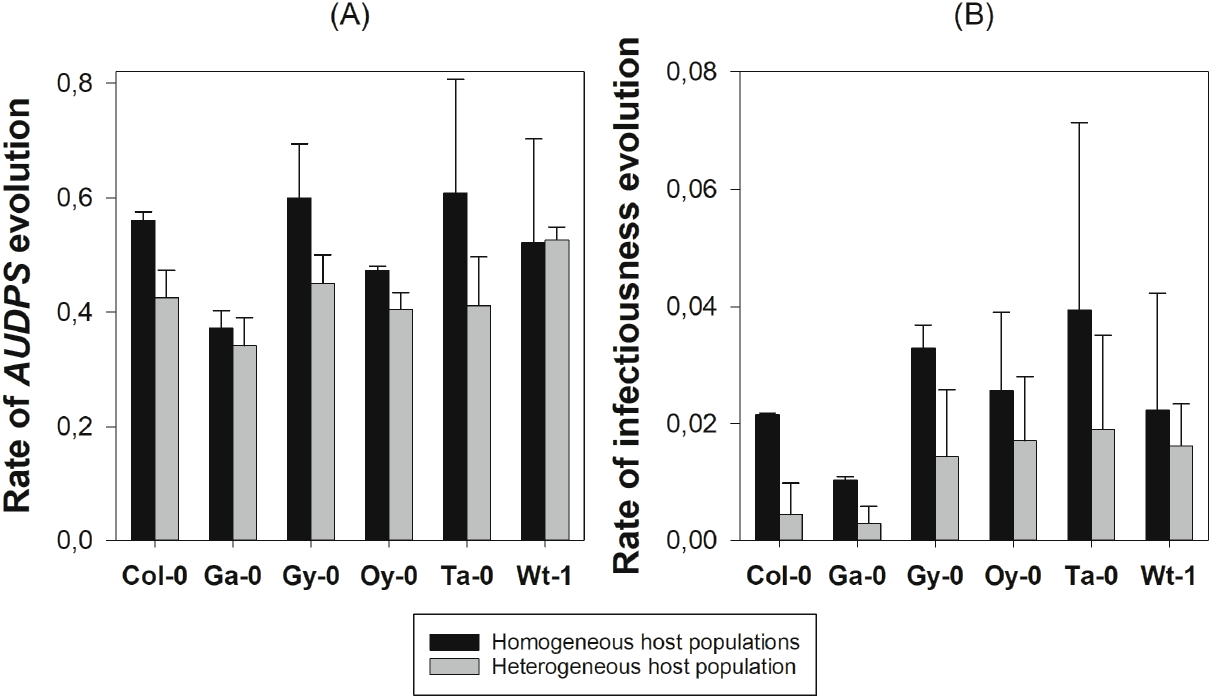
Mean rates of phenotypic evolution for the two traits studied, *AUDPS* and I. Rates of evolution were estimated from the ARIMA(1,0,0) model described in section 2.6. Error bars represent ±1 SEM.

**Table 2.** GLM analyses of the of the rates of evolution of *AUDPS* and *I* data (Fig. 5). See section 2.6 for a description of the model equation and parameters. 1 – *β* is the power of the corresponding test.

| Source of variation<sup>1</sup> | Rate of evolution ($\beta^Y$) | $\chi^2$ | d.f. | $p$ | $\eta_p^2$ | $1 - \beta$ |
| --- | --- | --- | --- | --- | --- | --- |
| *Intersection* ($\overline{\beta^Y}$) | *AUDPS* | 80.518 | 1 | < 0.001 | 0.965 | 1 |
|  | *I* | 25.879 | 1 | < 0.001 | 0.660 | 0.993 |
| *D* | *AUDPS* | 5.936 | 1 | 0.015 | 0.219 | 0.393 |
|  | *I* | 4.979 | 1 | 0.026 | 0.187 | 0.334 |
| *E* | *AUDPS* | 8.835 | 5 | 0.116 | 0.308 | 0.259 |
|  | *I* | 6.176 | 5 | 0.289 | 0.227 | 0.179 |
| *D*×*E* | *AUDPS* | 3.378 | 5 | 0.642 | 0.131 | 0.110 |
|  | *I* | 1.071 | 5 | 0.975 | 0.044 | 0.067 |

<sup>1</sup>Model factors: population structure $\equiv D \in$ (homogeneous, heterogeneous), host ecotype $\equiv E \in$ (Col-0, Ga-0, Gy-0, Oy-0, Ta-0, Wt-1).

Secondly, a similar result has been observed for the rates of *I* evolution (Fig. 5B): rates of infectiousness evolution are faster in the homogeneous populations than in the heterogeneous ones. Again, differences among population structures are significant (Table 2; *p* = 0.026) and of large effect 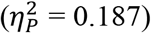.

For both traits, no significant differences in rates of evolution exist between ecotypes (*E*) nor for the interaction between ecotypes and population structure (*D* × *E*) (Table 2), with the power of the tests being too low (1 – *β* ≤ 0.259) as to fully discard we are not accepting a false null hypothesis of no effects. Low statistical power must clearly be due to the small sample size used (only two experimental blocks).

As a conclusion for this second section, evolution of virulence and infectiousness took place at a faster pace in genetically homogeneous host populations than in heterogeneous populations.

### 3.3. Comparison of infection matrices

Finally, we evaluated the degree of specificity of adaptation to the six ecotypes for all the evolved lineages. To build up infection matrices, TuMV lineages evolved in each of the ecotypes, or isolated from each ecotype in the case of the two lineages evolved in the heterogeneous host population, were inoculated into each one of the six ecotypes and *AUDPS* was estimated as described above. Fig. 6 shows a binary representation of the infection matrices estimated (averaging across experiments) for the two host population structures. Black squares represent host-virus combinations in which virulence was equal or greater than for the value observed for the viral lineage in its corresponding local host (host of isolation in the case of the heterogenous host population). White squares represent less virulent combinations. The first row in the matrices correspond to the most generalist TuMV lineage, in this case those evolved in Gy-0, whereas the last row represents the most specialist TuMV lineage, here those evolved in Oy-0. This observation is consistent for both matrices. Likewise, both matrices are also consistent regarding the rank order in susceptibility to infection of the different ecotypes: Oy-0 is the most susceptible ecotype, being infected by almost all viral lineages with similar virulence (exception being the virus isolated from Col-0 in the heterogenous host population) and Gy-0 being the most resistant ecotype, infected with high virulence only by the lineages evolved in or isolated from Gy-0. In other words, this suggests that more permissive ecotypes select for less virulent viruses while more restrictive ecotypes select for more virulent ones.

**Figure 6.**
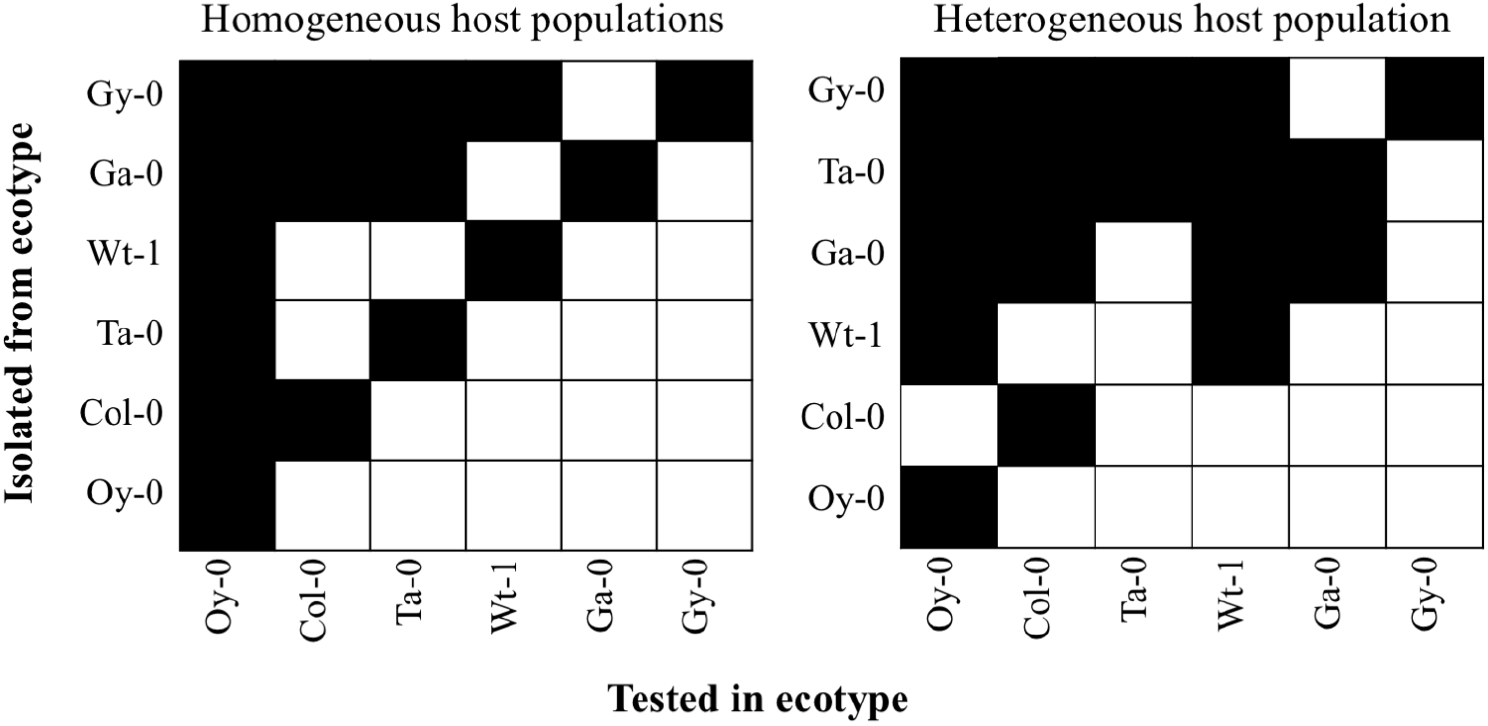
Infection matrices obtained from the virulence data (mean *AUDPS* values of the two experiments). Black cells represent cases in which virulence was equal or greater than the value estimated for the corresponding viral lineage on its local host ecotype. In the case of the heterogeneous population, it corresponds to the value estimated on the ecotype from which the virus was isolated in the last evolution passage. As described in section 3.2, in each matrix evolved viruses are ordered from the most virulent (upper row) to the less virulent (bottom row) and host ecotypes from the most sensitive (most left column) to the most resistant (rightest column).

To further test this hypothesis, we evaluated the nestedness and modularity of the two matrices. The infection matrix estimated for the homogeneous host populations shows a significant nestedness (Fig. 6, left; *p* = 0.035). This suggests that virus evolution in a single host genotype selects for a gene-for-gene interaction mechanism. This model of host-virus interaction is fully compatible with the above hypothesis. However, the matrix estimated for the heterogenous population did not show significant nestedness (Fig. 6, right; *p* = 0.149), suggesting that gene-for-gene interactions have not being selected under this ecological situation. Indeed, odds ratios indicate that the left matrix in Fig. 6 is 3.16% more nested that the right one.

Both matrices show significant modularity (*p* = 0.022 for the homogeneous and *p* = 0.012 for the heterogeneous host populations), though the modularity in the heterogeneous host population matrix is slightly larger (odds ratio: 0.24%). A module is a group of viruses and hosts for which the viruses in the set are more likely to be virulent in these hosts that to any other host outside the group, and that hosts in the group are more likely to be infected with similar virulence by viruses from within the group. Such modules suggest common selective constraints imposed by the hosts and similar evolutionary solutions found by the viruses.

Finally, we sought to quantify the degree of specialization in the host ecotype – viral lineage, or partner diversity. To this end, we applied Blüthgen et al. (2006) *d*’ index. For the infection matrix estimated for viruses evolved in the homogeneous host populations (Fig. 6 left), *d*’ = 0.00207, while the for the matrix estimated for viruses evolved in the heterogeneous host population (Fig. 6 right), *d*’ = 0.00061. Therefore, 3.39 times greater specialization evolved when the virus was facing a single host genotype during the evolution experiment.

## 4. Discussion

### 4.1. Host population structure and the evolution of specialist and generalist viral strategies

Most plant viruses are generalists capable of infecting more than one host species and thus generalism *versus* specialism should be properly defined on the basis of variance in infectiousness across different hosts (Leggett et al. 2013). In this sense, a generalist virus will be characterized by a low variance in infectiousness over different host genotype and/or species; by contrast a specialist virus will have a high variance. It is logical to expect that virus evolution should proceed faster in homogenous than in heterogeneous host environments because viruses with a narrower host range (specialist) have higher probabilities of fixing beneficial alleles, taking less time to do so (Gavrilets and Gibson 2002; Whitlock 2002, 2003; Whitlock and Gomulkiewicz 2005; Papaïx et al. 2013). Consequently, viral species or genotypes with broader host ranges (generalists) must have slower rates of evolutionary response (Bennett et al. 1992; Fry 1993; Whitlock 1996; Kassen and Bell 1998; Kassen 2002). If evolution occurs in too many hosts, then the total selection for any particular host-specific trait would not be as effective, and viruses specializing into a single host would then evolve faster and outcompete their generalist counterparts. As a result of this faster evolutionary rate, specialist viruses may persist longer in time in a constant host landscape. In this study, we have directly challenged this hypothesis using experimental evolution. Lineages of TuMV, a prototypical RNA virus of the picorna-like superfamily (*sensu* Koonin et al. 2008), were evolved either in six alternative single host environments or in a genetically heterogeneous host environment composed by equal numbers of the same six host genotypes. Giving support to the above hypothesis, we observed that rates of phenotypic evolution were significantly faster for lineages evolved in the homogeneous host populations than in the heterogeneous one.

Pfenning (2001) made the distinction between polymorphism and polyphenism as causes of pathogens’ generalism. At the one hand, polymorphism means that different strains of pathogens evolve specialized virulence strategies in different host genotypes. At the other hand, polyphenism means that pathogens facultatively express alternative virulence strategies depending on the host phenotypes. In case of viruses with compacted genomes and multifunctional proteins, polymorphisms seem a more plausible explanation, though we cannot rule out polyphenism. In this sense, highly polymorphic viral populations would behave as generalists owed to the diversity of specialists they contain.

### 4.2. Host population structure, selection of recognition mechanisms and evolution of virulence

We have also characterized the evolution of two-virulence related traits, *AUDPS* and infectiousness. According to theoretical predictions (Haraguchi and Sasaki 2000; Regoes et al. 2000; Gandon 2004; Lively 2010; Moreno-Gámez et al. 2013), one should expect to observe viruses to evolve toward mild infections in genetically homogeneous host populations, as a more efficient exploitation strategy that maximizes transmission. By contrast, host heterogeneity is postulated to select for increased virulence (Ganusov et al. 2002). Indeed, in the absence of trade-offs between the virulence in the different host ecotypes, it is expected that virulence increases without bounds (Regoes et al. 2000). Our results also provide support to this hypothesis. TuMV lineages evolved in homogenous host populations are, on average, less virulent on their local hosts than those lineages evolved in the maximally heterogeneous host population. However, a note of caution must be added here: the aforementioned models assume a trade-off between virulence and transmission rates because more virulent viruses will reduce their host’s lifespan or production of viable progeny. In our experiments, this trade-off has been broken since transmission rate is controlled by us and, therefore, virulence can increase without paying a cost. Therefore, our results are *consistent* with the expectation that mild infections are selected in genetically homogenous host populations, though the precise mechanism does not rely in the transmission-virulence trade-off and needs to be explored in future work.

Genetic variability in host’s susceptibility to infection and the evolution of infectiousness and virulence of parasites has been well documented in animals and plants (see Schmid-Hempel and Koella (1994) and Parrat et al. (2016) for reviews). Resistance is not always achieved against all parasite genotypes but results from the specific interaction between the genotypes of the host and the pathogen in agreement with the so-called gene-for-gene relationship. This mechanism proposes that for each locus determining resistance in the plant host, there is a corresponding locus for virulence in the virus. This model predicts that infection matrices must show significant nestedness: the most susceptible host carrying few resistance genes will be successfully infected by every virus, while the host genotypes carrying large number of resistance loci will be infected by very few viral genotypes carrying more virulence loci (Weitz et al. 2013). To test whether the gene-for-gene relationship has evolved in our experiment, we evaluated the structure of the infection matrices. In agreement with the gene-for-gene model, we found that the infection matrix for the TuMV lineages evolved in homogeneous hosts was significantly nested, with viruses evolved in the most restrictive host Gy-0 being also the most generalist ones able of infecting all alternative ecotypes equally well. By contrast, viruses evolved in the most permissive host Oy-0 were the most specialized ones, infecting alternative hosts with less efficiency. Interestingly, the lineages evolved in the heterogeneous host population did not generate a nested matrix, suggesting that fluctuating selection avoided the fixation of beneficial alleles in all required virulence loci. It is also noteworthy mentioning that, regardless whether or not nested, both infection matrices were significantly modular, suggesting that the mechanism underlying TuMV-A. *thaliana* interaction is far more complex than expected under a simple gene-for-gene model. Two possible mechanisms will contribute to create modularity: firstly, plant ecotypes sharing alleles in given sets of defense-response genes will likely impose similar selective constraints to the virus. Secondly, negative pleiotropic effect of mutations in small and compacted RNA genomes limits the number of alternative evolutionary pathways.

### 4.3. Are our findings in agreement with those from other previous evolution experiments?

We would like to compare our findings with those from studies with small RNA and large DNA viruses. First, Cuevas et al. (2003) simulated *in vitro* the migration of *Vesicular stomatitis virus* (VSV) between three alternative cell types, and observed that migration rate had a negative impact on the rate of fitness improvement on each cell type. With no migration, viruses evolved as specialists with high fitness in their local host, while at 50% migration rate per generation VSV evolved as a lower fitness generalist metapopulation. This result is apparently at odds with our findings and with theoretical expectations. However, it is worth noticing that Cuevas et al. (2003) did not measure virulence but fitness in competition against a common reference strain. Since fitness and virulence are not perfectly correlated traits in cell cultures for VSV (Furió et al. 2012), this may account for the discrepancy.

Second, Hillung et al. (2014) evolved TEV, another member of the *Potyvirus* genus, in five different ecotypes of *A. thaliana* in a set up similar to our homogeneous host populations treatment. In full agreement with our results, the infection matrix of the evolved TEV genotypes on the five ecotypes was significantly nested and thus compatible with a gene-for-gene relationship.

Third, Boots and Mealor (2007) evolved lineage of the large DNA betabaculovirus PiGV in lines of its host lepidopteran at varying viscosity of the food media in which larvae grew. Increasing viscosity decreased the mobility of the larvae and thus created stronger spatial structure. Viruses evolved in spatially structured host populations evolved to be less virulent than those maintained in well mixed host populations, matching our findings.

Finally, Berngruber et al. (2015) showed that a latent λ phage won competitions against a virulent one in a spatially-structured bacterial host populations but lost in well mixed ones. This result agrees with our observation that evolution in isolated homogeneous host populations selected for less virulent TuMV while evolution in well mixed heterogeneous population selected for more virulent virus.

### 4.4. Are our results compatible with field observations?

There is a number of field studies performed with plant viruses whose results are fully compatible with those from our experimental approach. We are just mentioning a few. Malpica et al. (2006) analyzed the prevalence of five plant viruses on 21 wild plant species and found that specialization was the most successful viral strategy, with highly selective viruses (specialist) being the most prevalent ones. In a recent follow up study, McLeish et al. (2017) have applied ecological networks theory to explore the association between plant host diversity and virus spatial distributions. Two important observations were made: (i) an association between host abundance and viral prevalence and (ii) most prevalent viruses behave as facultative generalists, meaning that they have the widest host range breadth but can narrow it to the most permissive host without paying a fitness cost.

Rodelo-Urrego et al. (2013) explored the association between the population structure of *Capsicum annuum glabriusculum* and the prevalence of pepper golden mosaic and pepper huasteco yellow vein begomoviruses, founding that landscape heterogeneity affected the epidemiology and genetic structure of the begomoviruses. When host population diversity was removed by human intervention, the prevalence of the viruses increased. Higher levels of anthropization and loss of plant biodiversity (i.e., similar to our genetically homogenous host populations) resulted in an increase in the genetic diversity of the begomoviruses (Rodelo-Urrego et al. 2015).

### 4.5. Concluding remarks

Ecological factors, such as the spatial distribution of host genotypes, is fundamental to understand the evolutionary and epidemiological dynamics of infectious diseases. The role of spatial host structure has attracted quite a lot of attention from theoreticians that made a number of interesting testable predictions. Unfortunately, many of these still remain untested. Here we have tried to provide experimental evidences for a plant-virus pathosystem to test the hypothesis that host population homogeneity would promote fast local adaptation and low virulence of the virus, whereas the presence of host genetic heterogeneity for susceptibility to infection will slow down the rate of viral evolution and favor more virulent viruses. Supporting these predictions, we found faster TuMV adaptation to homogeneous than to heterogeneous *A. thaliana* experimental populations. However, viruses evolved in heterogeneous host populations were more pathogenic and infectious than viruses evolved in the homogeneous population. Furthermore, the viruses evolved in homogeneous populations showed stronger signatures of local specialization than viruses evolved in heterogeneous populations. These results illustrate how the genetic diversity of hosts in an experimental ecosystem favors the evolution of virulence of a pathogen and may help agronomists to handle crops in ways that will minimize the rise and spread of virulent viral strains.

## Supplementary data

Supplementary data are available at *Virus Evolution* online.

Raw data, including numbers of symptomatic plants, computation of *AUDPS* and infectiousness values at each passage and experiment are available at LabArchives under doi: 10.25833/q8y2-f433.

## Acknowledgements

We thank Francisca de la Iglesia for continuous excellent technical support. Work was supported by Spain’s Agencia Estatal de Investigación – FEDER grant BFU2015-65037-P and Generalitat Valenciana grant GRISOLIA/2018/005 to S.F.E. R.G. was supported by Spain’s Agencia Estatal de Investigación pre-doctoral contract BES-2016-077078.

---

**Supplementary Table 1.** Results of the univariate MANOVAs. Model equation fitted to each phenotypic variable is the same as for the multivariate analyses (section 2.7).

| Source | Variable | SS | df | MS | F | P | $\eta_p^2$ | $1 - \beta$ |
| --- | --- | --- | --- | --- | --- | --- | --- | --- |
| *Intersection* | *AUDPS* | 1777.003 | 1 | 1777.003 | 814.700 | < 0.001 | 0.755 | 1 |
|  | *f* | 44.188 | 1 | 44.188 | 302.233 | < 0.001 | 0.534 | 1 |
| *t* | *AUDPS* | 978.630 | 1 | 978.630 | 448.671 | < 0.001 | 0.630 | 1 |
|  | *f* | 11.722 | 1 | 11.722 | 80.175 | < 0.001 | 0.233 | 1 |
| *B* | *AUDPS* | 152.964 | 1 | 152.964 | 70.129 | < 0.001 | 0.210 | 1 |
|  | *f* | 5.698 | 1 | 5.698 | 38.970 | < 0.001 | 0.129 | 1 |
| *B×t* | *AUDPS* | 0.448 | 1 | 0.448 | 0.206 | 0.651 | 0.001 | 0.074 |
|  | *f* | 2.547 | 1 | 2.547 | 17.420 | < 0.001 | 0.062 | 0.986 |
| *D* | *AUDPS* | 25.107 | 1 | 25.107 | 11.511 | 0.001 | 0.042 | 0.922 |
|  | *f* | 2.083 | 1 | 2.083 | 14.244 | < 0.001 | 0.051 | 0.964 |
| *D×t* | *AUDPS* | 9.070 | 1 | 9.070 | 4.158 | 0.042 | 0.016 | 0.529 |
|  | *f* | 1.100 | 1 | 1.100 | 7.523 | 0.007 | 0.028 | 0.780 |
| *E* | *AUDPS* | 175.155 | 5 | 35.031 | 16.061 | < 0.001 | 0.233 | 1 |
|  | *f* | 3.608 | 5 | 0.722 | 4.935 | < 0.001 | 0.085 | 0.982 |
| *E×t* | *AUDPS* | 14.695 | 5 | 2.939 | 1.347 | 0.245 | 0.025 | 0.474 |
|  | *f* | 1.168 | 5 | 0.234 | 1.598 | 0.161 | 0.029 | 0.553 |
| *B×D* | *AUDPS* | 4.236 | 1 | 4.236 | 1.942 | 0.165 | 0.007 | 0.284 |
|  | *f* | 0.128 | 1 | 0.128 | 0.874 | 0.351 | 0.003 | 0.154 |
| *B×D×t* | *AUDPS* | 8.599 | 1 | 8.599 | 3.942 | 0.048 | 0.015 | 0.507 |
|  | *f* | 0.288 | 1 | 0.288 | 1.970 | 0.162 | 0.007 | 0.288 |
| *B×E* | *AUDPS* | 15.355 | 5 | 3.071 | 1.408 | 0.222 | 0.026 | 0.494 |
|  | *f* | 2.863 | 5 | 0.573 | 3.916 | 0.002 | 0.069 | 0.943 |
| *B×E×t* | *AUDPS* | 16.678 | 5 | 3.336 | 1.529 | 0.181 | 0.028 | 0.532 |
|  | *f* | 1.514 | 5 | 0.303 | 2.072 | 0.069 | 0.038 | 0.685 |
| *D×E* | *AUDPS* | 3.512 | 5 | 0.702 | 0.322 | 0.900 | 0.006 | 0.131 |
|  | *f* | 0.179 | 5 | 0.036 | 0.245 | 0.942 | 0.005 | 0.110 |
| $D \times E \times t$ | $AUDPS$ | 5.351 | 5 | 1.070 | 0.491 | 0.783 | 0.009 | 0.183 |
| | $f$ | 0.175 | 5 | 0.035 | 0.239 | 0.945 | 0.005 | 0.108 |
| $B \times D \times E$ | $AUDPS$ | 6.331 | 5 | 1.266 | 0.581 | 0.715 | 0.011 | 0.212 |
| | $f$ | 0.349 | 5 | 0.070 | 0.478 | 0.793 | 0.009 | 0.179 |
| $B \times D \times E \times t$ | $AUDPS$ | 8.256 | 5 | 1.651 | 0.757 | 0.582 | 0.014 | 0.271 |
| | $f$ | 0.520 | 5 | 0.104 | 0.711 | 0.616 | 0.013 | 0.255 |
| Error | $AUDPS$ | 575.830 | 264 | 2.181 | | | | |
| | $f$ | 38.598 | 264 | 0.146 | | | | |

**Supplementary Table 2.**
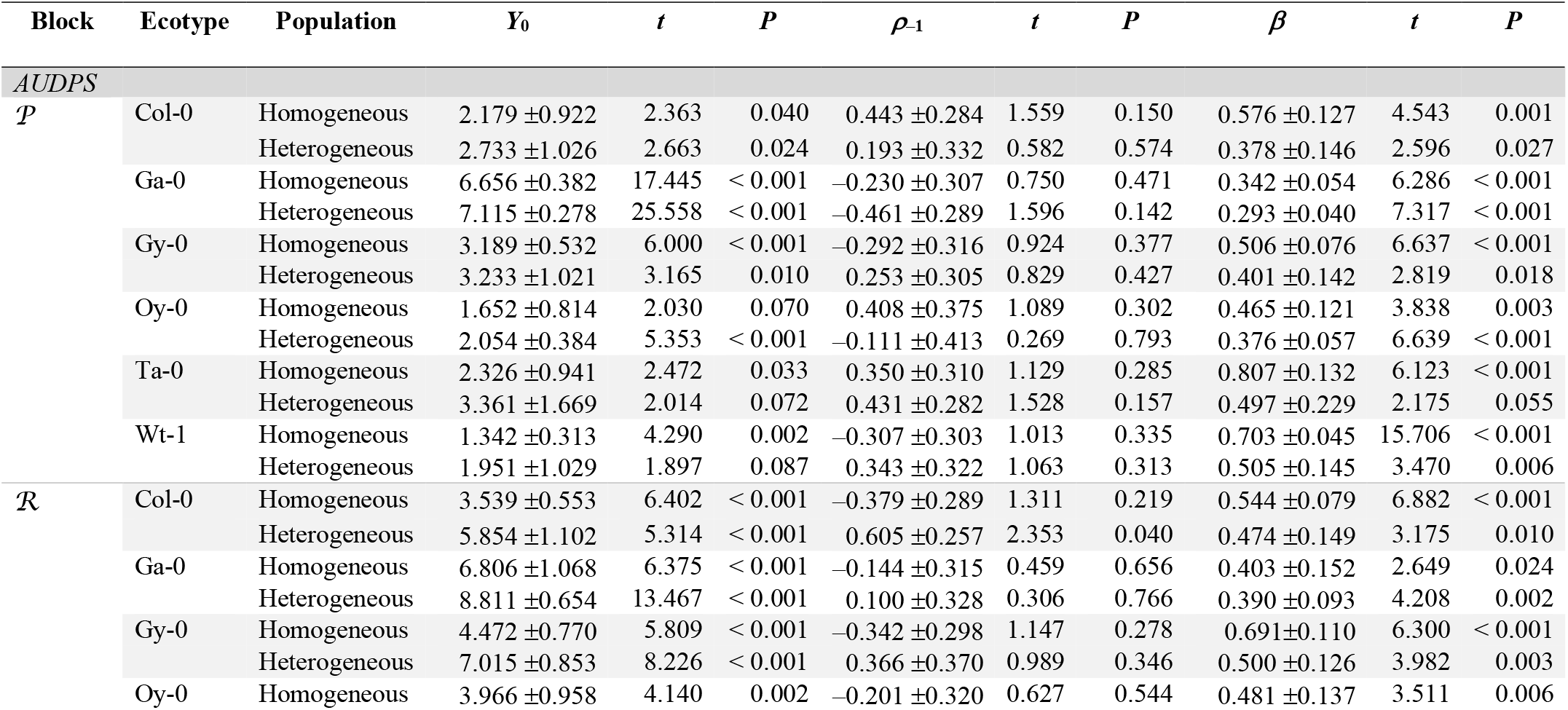
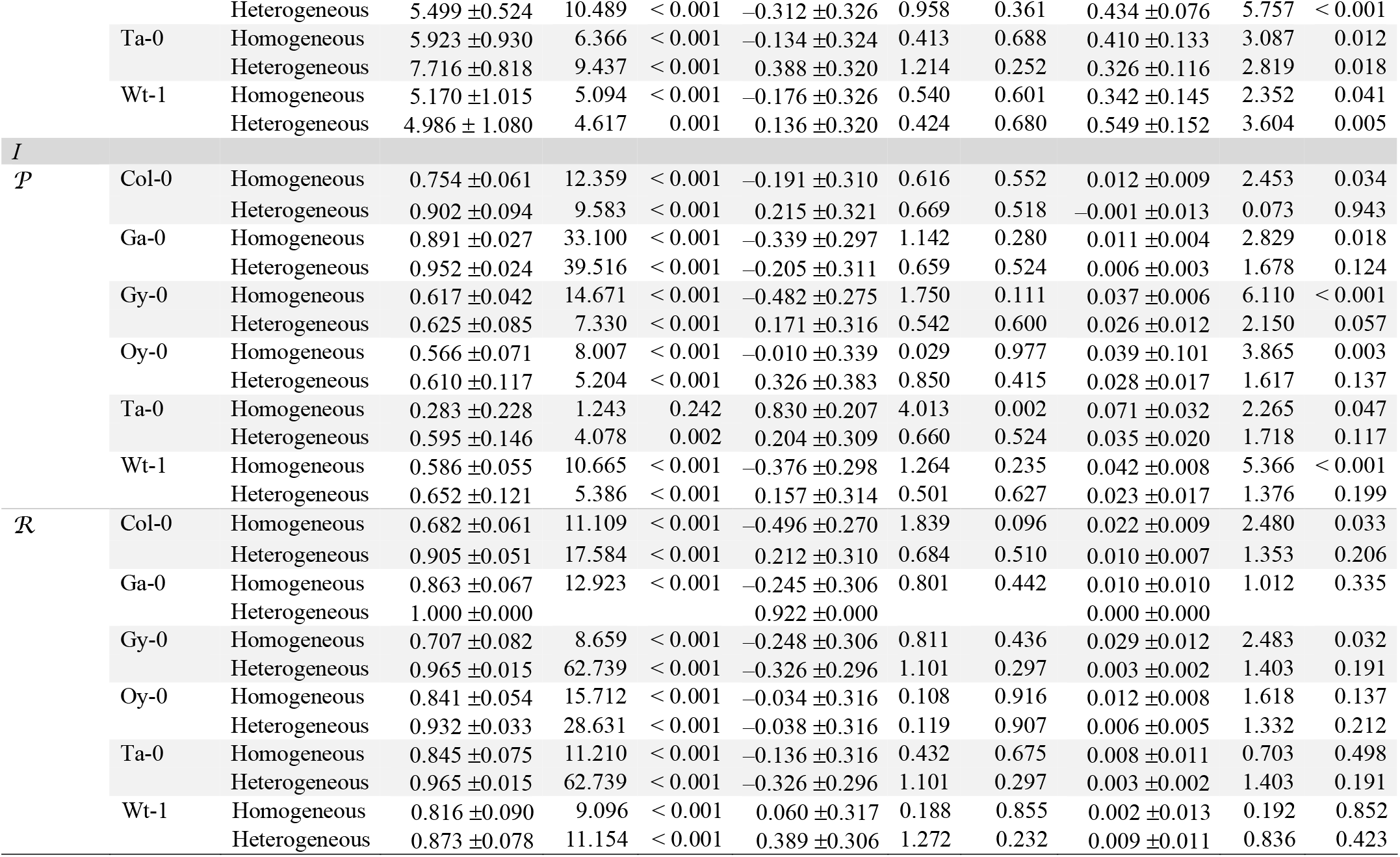
Results of the fitting of the ARIMA(1,0,0) model 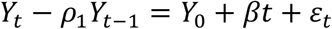 to the *AUDPS* and *I* time series data. Error represent ±1 SEM. See section 2.6 for definition of each model parameter. Statistical significance of each parameter is estimated using a *t*-test (10 d.f.).

## References

Altizer, S. et al. (2006) “Seasonality and the dynamics of infectious diseases”. Ecology Letters, 9: 467–484.

Anttila, J. et al. (2015) “Environmental variation generates environmental opportunist pathogen oubreaks”. PLoS ONE, 10: e0145511.

Bascompte, J. et al. (2003) “The nested assembly of plant-animal mutualistic networks”. Proceedings of the National Academy of Sciences of the USA, 100: 9383–9387.

Baylis, M. (2017) “Potential impact of climate change on emerging vector-borne and other infections in the UK”. Environmental Health, 16: 112.

Bennett, A. F., Lenski, R. E., Mittler, J. E. (1992) “Evolutionary adaptation to temperature. I. Fitness responses of *Escherichia coli* to changes in its thermal environment”. Evolution, 46: 16–30.

Berngruber, T. W., Lion, S., Gandon, S. (2015) “Spatial structure, transmission modes and the evolution of viral exploitation strategies”. PLoS Pathogens, 11: e1004810.

Blüthgen, N., Menzel, F., Blüttgen N. (2006) “Measuring specialization in species interaction networks”. BMC Ecology, 6: 12.

Boots, M., Mealor, M. (2007) “Local interactions select for lower pathogen infectivity”. Science, 315: 1284–1286.

Boots, M., Sasaki, A. (1999) “‘Small worlds’ and the evolution of virulence: infection occurs locally and at a distance”. Proceedings of the Royal Society B: Biological Sciences, 266: 1933–1938.

Boots, M., Sasaki, A. (2002) “Parasite-driven extinction in spatially explicit host-parasite systems”. American Naturalist, 159: 706–713.

Boyes, D. C. et al. (2001) “Growth stage-based phenotypic analysis of Arabidopsis: a model for high throughput functional genomics in plants”. Plant Cell, 13: 1499–1510.

Brockhurst, M. A., Buckling, A., Rainey, P. B. (2006) “Spatial heterogeneity and the stability of host-parasite coexistence”. Journal of Evolutionary Biology, 19: 374–379.

Brown, J. K. M., Tellier, A. (2011) “Plant-parasite coevolution: bridging the gap between genetics and ecology”. Annual Review of Phytopathology, 49: 345–367.

Chabas, H. et al. (2018) “Evolutionary emergence of infectious diseases in heterogeneous host populations”. PLoS Biology, 16: e2006738.

Chen, C. C. et al. (2003) “Identification of *Turnip mosaic virus* isolates causing yellow stripe and spot on calla lily”. Plant Disease, 87: 901–905.

Comins, H. N., Hassell, M. P., May, R. M. (1992) “The spatial dynamics of host parasitoid systems”. Journal of Animal Ecology, 61: 735–748.

Cuevas, J. M., Moya, A., Elena, S. F. (2003) “Evolution of RNA virus in spatially structured heterogeneous environments”. Journal of Evolutionary Biology, 16: 456–466.

Di Giallonardo, F., Holmes, E. C. (2015) “Viral biocontrol: Grand experiments in disease emergence and evolution”. Trends in Microbiology, 23: 83–90.

Dormann CF, Gruber B, Fruend J. (2008) “Introducing the bipartite package: analyzing ecological networks”. R news, 8: 8–11.

Engering, A., Hogerwerf, L., Slingenbergh, J. (2013) “Pathogen-host-environment interplay and disease emergence”. Emerging Microbes and Infections, 2: e5.

Fry, J. D. (1993) “The evolution of host specialization: are trade-offs overrated?”. American Naturalist, 148: S84–S107.

Furió, V. et al. (2012) “Relationship between within-host finess and virulence in the *Vesicular stomatitis virus:* correlation with partial decoupling”. Journal of Virology, 86: 12228–12236.

Gandon, S. (2004) “Evolution of multihost parasites”. Evolution, 58: 455–469.

Gandon, S. et al. (1996) “Local adaptation and gene-for-gene coevolution in a metapopulation model”. Proceedings of the Royal Society B: Biological Sciences, 263: 1003–1009.

Gandon, S., Michalakis, Y. (2002) “Local adaptation, evolutionary potential and host-parasite coevolution: interactions between migration, mutation, population size and generation time”. Journal of Evolutionary Biology, 15: 451–462.

Ganusov, V. V., Bergstrom, C. T., Antia, R. (2002) “Within-host population dynamics and the evolution of microparasites in a heterogeneous host population”. Evolution, 56: 213–223.

Garrett, K. A. et al. (2006). “Climate change effects on plant disease: genomes to ecosystems”. Annual Review of Phytopathology, 44: 489–509.

Gavrilets, S., Gibson, N. (2002) “Fixation probability in a spatially heterogeneous environment”. Population Ecology, 44: 51–58.

Haraguchi, Y., Sasaki, A. (2000) “The evolution of parasite virulence and transmission rate in a spatially structured population”. Journal of Theoretical Biology, 203: 85–96.

Hillung, J. et al. (2014) “Experimental evolution of an emerging plant virus in host genotypes that differ in their susceptibility to infection”. Evolution, 68: 2467–2480.

Kassen, R. (2002) “The experimental evolution of specialists, generalists, and the maintenance of diversity”. Journal of Evolutionary Biology, 15: 173–190.

Kassen, R., Bell, G. (1998) “Experimental evolution in *Chlamydomonas.* IV. Selection in environments that vary through time at different scales”. Heredity, 80: 732–741.

Koonin, E. V. et al. (2008) “The Big Bang of picorna-like virus evolution antedates the radiation of eukaryotic supergroups”. Nature Review Microbiology, 6: 925–939.

Leggett, H. C. et al. (2013) “Generalism and the evolution of parasite virulence”. Trends in Ecology and Evolution, 10: 592–596.

Lively, C. M. (2010) “The effect of host genetic diversity on disease spread”. American Naturalist, 175: E149-E152.

Malpica, J. M. et al. (2006) “Association and host selectivity in multi-host pathogens”. PLoS ONE, 1: e41.

McLeish, M. et al. (2017) “Scale dependencies and generalism in host use shape virus prevalence”. Proceedings of the Royal Society B: Biological Sciences, 284: 20172066.

Moreno-Gámez, S., Stephan, W., Tellier, A. (2013) “Effect of disease prevalence and spatial heterogeneity on polymorphism maintenance in host-parasite interactions”. Plant Pathology, 62: S133–S141.

Morens, D. M., Fauci, A. S. (2013) “Emerging infectious diseases: threats to human health and global stability”. PLoS Pathogens, 9: e1003467.

Newman, M. E. J. (2006) “Modularity and community structure in networks”. Proceedings of the National Academy of Sciences of the USA, 103: 8577–8582.

Papaïx, J., et al. (2013) “Dynamics of adaptation in spatially heterogeneous metapopulations”. PLoS ONE, 8: e54697.

Parrat, S. R., Numminen, E., Laine, A. L. (2016) “Infectious disease dynamics in heterogeneous landscapes”. Annual Review of Ecology, Evolution and Systematics, 47: 283–306.

Pfenning, K. S. (2001) “Evolution of pathogen virulence: the role of variation in host phenotype”. Proceedings of the Royal Society B: Biological Sciences, 268: 755–760.

R Core Team. (2016) “R: A language and environment for statistical computing”. R Foundation for Statistical Computing, Vienna, Austria.

Regoes, R. R., Nowak, M. A., Bonhoeffer, S. (2000) “Evolution of virulence in a heterogeneous host population”. Evolution, 54: 64–71.

Rodelo-Urrego, M. et al. (2013) “Landscape heterogeneity shapes host-parasite interactions and results in apparent plant-virus codivergence”. Molecular Ecology, 22: 2325–2340.

Rodelo-Urrego, M., García-Arenal, F., Pagán, I. (2015) “The effect of ecosystem biodiversity on virus genetic diversity depends on virus species: a study of chiltepin-infecting begomovirus in Mexico”. Virus Evolution, 1: vev004.

Rodríguez, D. J., Torres-Sorando, L. (2001) “Models of infectious diseases in spatially heterogeneous environments”. Bulletin of Mathematical Biology, 63: 547–571.

Rosenthal, S. R. et al. (2015) “Redefining disease emergence to improve prioritization and macro-ecological analyses”. One Health, 1: 17–23.

Rubio, B. et al. (2019) “Genome wide association study reveals new loci involve in *Arabidopsis thaliana* and *Turnip mosaic virus* (TuMV) interactions in the field” New Phytologist, 221: 2026–2038.

Schmid-Hempel, P., Koella, J.C. (1994) “Variability and its implications for host-parasite interactions”. Parasitology Today, 10: 98–102.

Simko, I., Piepho, H.P. (2012) “The area under the disease progress stairs: calculation, advantage, and application”. Phytopathology, 102: 381–389.

Thrall, P. H., Burdon, J. J: (2003) “Evolution of virulence in a plant host-pathogen metapopulation”. Science, 299: 1735–1737.

Thrall, P. H. et al. (2012) “Rapid genetic change underpins antagonistic coevolution in a natural host-pathogen metapopulation”. Ecology Letters, 15: 425–435.

Tellier, A., Brown, J. K. (2011) “Spatial heterogeneity, frequency-dependent selection and polymorphism in host-parasite interactions”. BMC Evolutionary Biology, 1: 11.

Vurro, M., Bonciani, B., Vannacci, G. (2010) “Emerging infectious diseases of crop plants in developing countries: impact on agriculture and socio-economic consequences”. Food Security, 2: 113.

Weitz, J. S. et al. (2013) “Phage-bacteria infection networks”. Trends in Microbiology, 21: 82–91.

Whitlock, M. C. (1996) “The Red Queen beats the jack-of-all-trades: the limitations on the evolution of phenotypic plasticity and niche breadth”. American Naturalist, 148: S65–S77.

Whitlock, M. C. (2002) “Selection, load and inbreeding depression in a large metapopulation”. Genetics, 160: 1191–1202.

Whitlock, M. C. (2003) “Fixation probability and time in subdivided populations”. Genetics, 164: 767–779.

Whitlock, M. C., Gomulkiewicz, R. (2005) “Probability of fixation in a heterogeneous environment”. Genetics, 171: 1407–1417.

Woolhouse, M. E. J., Dye, C. (2001) “Preface: Population Biology of Emerging and Reemerging Pathogens”. Philosophical Transactions of the Royal Society B: Biological Sciences, 356: 981–982.

Yates, A., Antia, R., Regoes, R. R. (2006) “How do pathogen evolution and host heterogeneity interact in disease emergence?”. Proceedings of the Royal Society B: Biological Sciences, 273: 3075–3083.

